# Transcriptional adaptation influences buffering among essential paralog gene families

**DOI:** 10.64898/2026.09.24.754217

**Authors:** Siobhan O’Brien, Marissa H. Fujimoto, Amy R. Lowe, Alice H. Berger

## Abstract

Buffering among paralogous genes is a well-described phenomenon in which genetic redundancy provides genetic robustness. Shared function among paralogs allows for a compensation effect when one or more paralog family genes are inactivated. Considering the role of transcriptional adaptation in influencing gene expression among highly homologous genes, we sought to identify whether transcriptional adaptation has a role in paralog buffering. We performed Perturb-seq of 110 paralog gene knockouts and assessed compensatory gene expression effects among paralogs as well as at the whole transcriptome level. This analysis identified six paralog pairs that undergo a transcriptional adaptation following CRISPR-mediated loss of one paralog in the pair. Further, we found that the upregulation of *TIA1* following paralog *TIAL1* loss requires nonsense-mediated decay, and active nuclear import via the importin IPO8. These data provide a comprehensive evaluation of transcriptional adaptation and show that transcriptional adaptation is recurrent but not pervasive among paralogous human genes.

## Main

Buffering among paralogs is a well-documented phenomenon, first observed in yeast genetic screens^1^, and subsequently identified through a myriad of mammalian CRISPR screens^2–7^ and mouse genetic studies where loss-of-function of predicted essential genes had minor effects due to compensation^8^. Though feedback loops constitute one mechanism of paralog buffering^9,10^, more recent work has identified an RNA-degradation-dependent mechanism of transcription-based adaptation among homologous genes^11–13^. Initially identified through differential effects of morpholino and CRISPR knockouts in zebrafish, several groups have uncovered that degradation of mutant mRNA is required to upregulate adapting genes in a sequence-specific manner^11,13^, termed degradation-dependent transcriptional adaptation (TA). Further, the role of degradation-dependent TA in disease processes is slowly being uncovered, with recent work in Duchenne muscular dystrophy^14^ outlining its role in altered disease phenotypes. Recently, the role of antisense RNA in mediating the TA response has been reported, which may suggest a broad role of TA in regulating gene expression due to the production of antisense RNAs at many transcriptional start sites^12,15^. These studies show that TA can serve as a mechanism of gene regulation to protect against deleterious mutations and identify genes with sequence homology as capable of undergoing mutation-driven TA. Paralogous genes, which emerged through evolutionary gene duplication events, often preserve both sequence homology and some elements of gene function. In human cells, approximately two-thirds of genes belong to multi-gene paralog families. While several of the first genes identified to undergo TA (*Rela*, *Actg1*, *Actb*) are part of paralog families, to date there has been no systematic assessment of TA across human paralogs.

In this work, we applied high-throughput functional genomics methods to assess transcriptional adaptation among highly essential human paralog gene pairs to determine how frequently this phenomenon occurs, particularly among genes sharing high sequence homology. Further, we identify two sets of paralog families, *NONO-PSPC1-SFPQ* and *TIA1-TIAL1*, that undergo degradation-independent or degradation-dependent transcriptional adaptation respectively. We further verify the role of nonsense-mediated decay (NMD) in this process and identify active nuclear import through the importin protein IPO8 as involved in mediating degradation-dependent TA.

## Results

### Perturb-seq with target enrichment identifies transcriptional adaptation among paralogs

Considering the high likelihood of CRISPRko to generate mutant mRNA^16^, we performed a pooled CRISPRko screen followed by single-cell RNA sequencing (Perturb-seq^17,18^) to identify how often transcriptional adaptation is occurring among paralog genes. Genes were selected based on their enhanced dependence in dual-knockout CRISPR screens previously published in the lung adenocarcinoma cell line PC9^4^. We generated a single knockout CRISPR library (“sgPEN”), selecting 440 sgRNAs corresponding to 55 essential pairs (110 paralog genes), with an average genetic interaction (GI) score of-1, an indication of strong dependence on the pair **(Fig.1a)**. PC9 cells were subjected to Perturb-seq **(Fig. 1b)** following infection and selection with the sgPEN library. Efficient gene targeting was confirmed by dropout of sgRNAs targeting essential genes after 25 days **(Fig S1a)**. Following isolation, cDNA sequencing libraries were enriched for the 110 paralog gene targets through hybridization capture leveraging biotinylated DNA probes and subsequent amplification **(Fig. 1b)**. Just over 36,000 cells passed standard QC filtering metrics, representing an average library coverage of 308 per gene target and 73 cells per gRNA, with 108 final paralog genes adequately represented **(Fig. S1bc)**.

**Figure 1:**
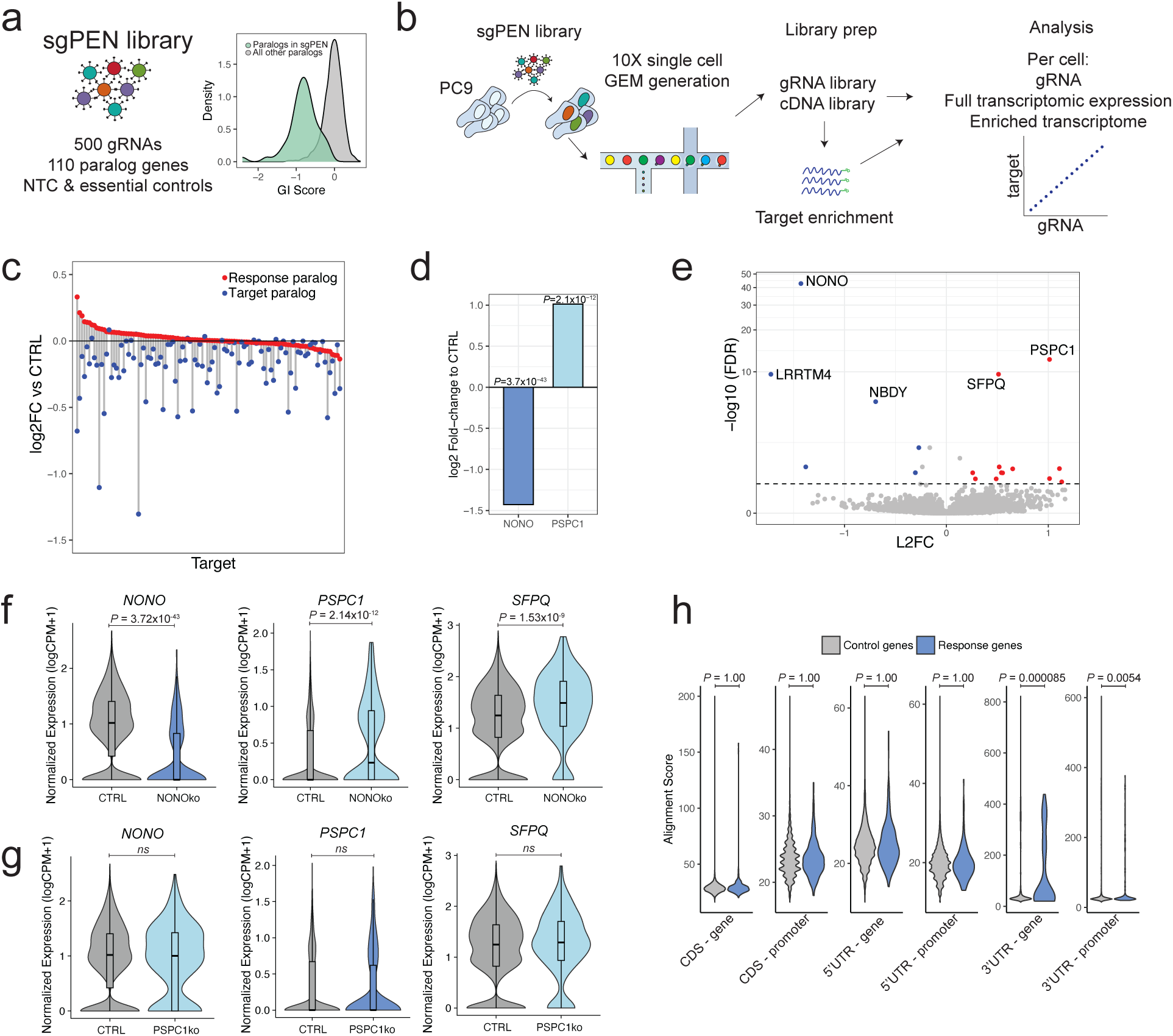
Transcriptional adaptation among paralog gene families a) Schematic of sgPEN CRISPR library (left) used in Perturb-seq primarily targets paralogs that are synthetic lethal in PC9 cells (right). NTC, non-targeting control. GI, genetic interaction. b) Experimental overview of paralog Perturb-seq assay using standard 10X protocols, with and without hybridization-capture of target paralogs (“enriched transcriptome”) c) Gene expression of each gene targeted by CRISPR knockout (“target”) vs paired “response” paralog expression across. Data shown is log_2_(fold change) of each gene in cells targeted with the gRNA of interest versus cells with non-targeting control gRNAs (CTRL) for 108 targeted genes in the whole transcriptome data set. d) Gene expression of *NONO* and *PSPC1* in cells containing sgRNAs targeting *NONO*, compared to cells containing control sgRNAs. P-values shown were generated by Wilcoxon rank-sum test and multiple hypothesis correction. Volcano plot of differential gene expression results comparing cells containing sgRNAs targeting *NONO* to cells containing control sgRNAs, points coloured by L2FC <-0.5 or > 0.5, FDR< 0.05. e) Gene expression of *NONO, PSPC1,* and *SFPQ* in *NONO*ko cells. P-values shown were generated by Wilcoxon rank-sum test and multiple hypothesis correction. f) Gene expression of *NONO, PSPC1,* and *SFPQ* mRNA expression in *PSPC1*ko cells. Wilcoxon rank-sum p-values were calculated and adjusted for multiple hypothesis testing. g) Sequence alignment scores between control genes and response genes, based on genomic features. P-values are adjusted Welch’s t-test.

### Whole transcriptome analysis uncovers adaptation signatures among higher-order paralogs

We first analyzed the whole transcriptome Perturb-seq libraries to provide a broader survey of transcriptional adaptation. Many paralog genes were adequately covered in the whole transcriptome data **(Fig. S1bc)**, and as expected, target genes showed knockout **(Fig. 1c)**.

Following differential gene expression analysis, only one pair was found to be significantly upregulated in response to target gene loss: *NONO* and *PSPC1* **(Fig. 1d)**. We next sought to understand whether other genes outside the initial pairings were upregulated in response to target paralog loss. All response genes with significant differential gene expression following *NONO* knockout were assessed; in addition to the upregulation of *PSPC1*, a third paralog in the family, *SFPQ*, was also upregulated **(Fig 1e)**. Cells with *NONO* knockout showed robust loss of *NONO* expression and upregulation of both *PSPC1* and *SFPQ* **(Fig. 1f)** whereas *PSPC1* knockout did not result in *NONO* or *SFPQ* upregulation **(Fig. 1g)**.

Previous reports of degradation-dependent TA describe a mechanism by which other sequence-similar genes, including but not restricted to paralogs, are upregulated in response to degradation of the target gene^11,13^. To assess whether this was occurring in our dataset, we first identified significantly upregulated response genes across all target genes in the dataset **(Fig. S1d)**. Next, sequence alignment scores for all target paralogs and their corresponding response genes were generated using pairwise alignment between MANE-select mRNA isoforms of target genes and gene sequences of response genes. mRNA sequences were split into 5’ untranslated region (UTR), CDS, and 3’ UTR fragments, while DNA sequences were divided between promoter regions (+2kb from transcriptional start site) and annotated gene sequence **(Fig. S1e)**. Alignment scores were then compared between response genes and a subset of genes that had no change in gene expression following target loss. Strikingly, 3’ UTRs of response genes had significantly higher alignment scores when compared to target gene promoters, and target gene coding sequences **(Fig. 1h)**. Despite lower selective pressure at 3’ UTRs than coding regions^19^, purifying selection does occur in this regulatory-rich region of mRNA, indicating the presence of important functional sequences in these regions. The mechanism behind why some 3’ UTR regions show patterns of homology to response genes at both the promoter and gene coding sequence of response genes is unclear, though homology between 3’ UTRs of homologous genes has been described^20^, and repetitive sequences could be contributing in this region. Despite this, it is well known that miRNAs target the 3’ regions of mRNAs to induce degradation and suppression of expression; perhaps some of the shared homology of miRNA target sites may underlie this homology. Recent work identifying TA trigger sequences in a mouse model identified several regions of sequence that can trigger upregulation of response genes, primarily driven by coding-coding homology, but 3’ UTR regions were also identified as TA triggers^12^.

### Target enrichment identifies an extended set of paralogs undergoing transcriptional adaptation

We next analyzed the paralog target-enriched transcriptome dataset to better assess both knockout efficiency and subsequent response gene upregulation focused specifically on our paralog gene set. Target enrichment successfully increased cell coverage for nearly all genes **(Fig. 2a)** with UMI values correlated to non-enriched samples (Pearson r= 0.97, p-value <2.2e6) **(Fig. S1f)**, and an overall read enrichment of 32X overall **(Fig. S1g)**.

**Figure 2:**
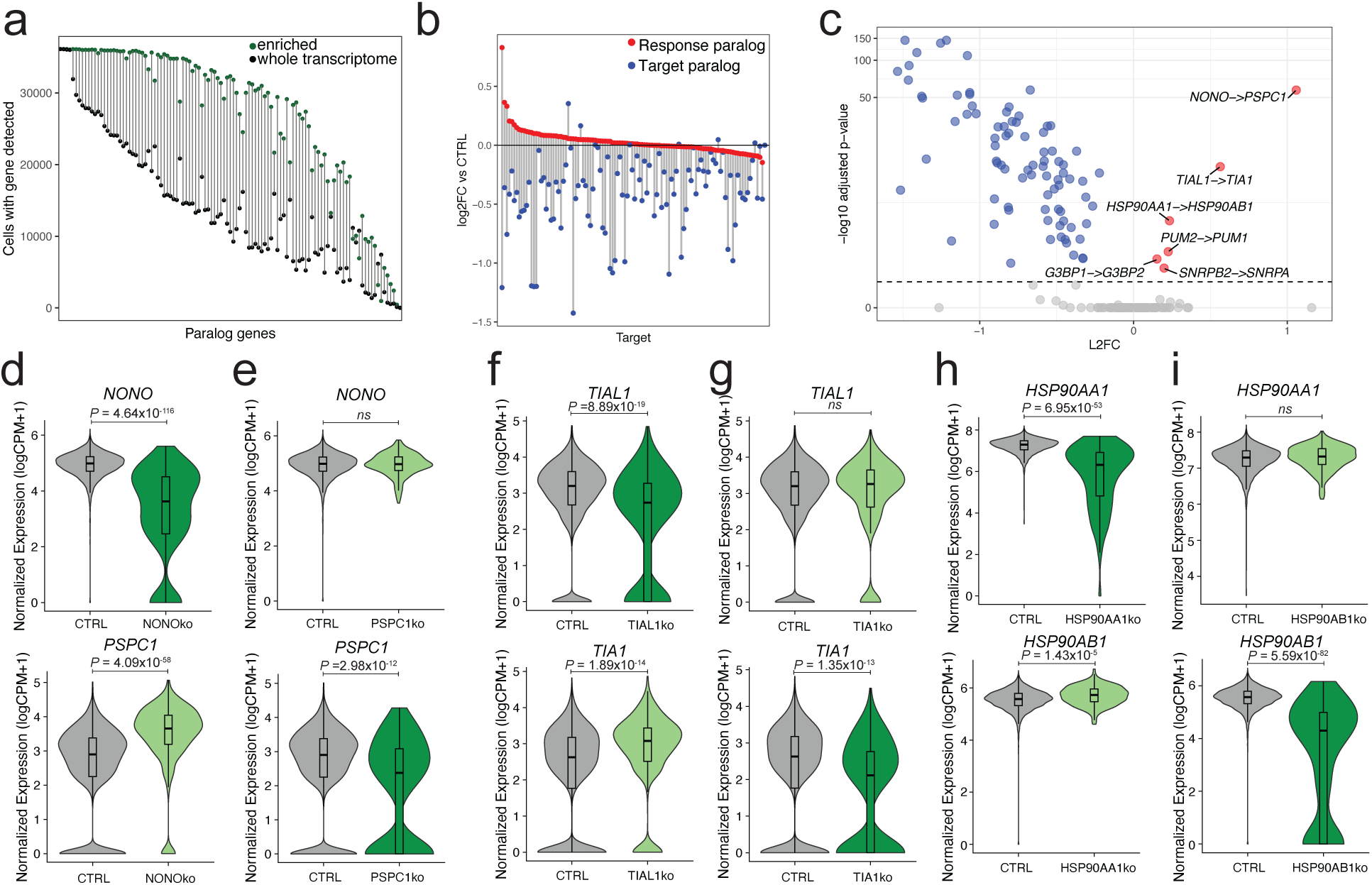
Perturb-seq with target enrichment identifies an expanded set of paralog pairs undergoing transcriptional adaptation a) Increased cell counts in enriched versus whole transcriptome datasets b) Gene expression of each gene targeted by CRISPR knockout (“target”) vs paired “response” paralog expression across. Data shown is log_2_(fold change) of each gene in cells targeted with the gRNA of interest versus cells with non-targeting control gRNAs (CTRL) for 108 targeted genes in the enriched data set. c) Target and response gene expression following pooled CRISPR knockout of paralogs, presented as “*target->response*” gene. A dashed line indicates Wilcoxon rank-sum test, adjusted p-value <0.05. d) Gene expression of *NONO* and *PSPC1* in *NONO*ko cells in the enriched Perturb-seq data. e) Gene expression of *NONO* and *PSPC1* in *PSPC1*ko cells in the enriched Perturb-seq data. f) Gene expression of *TIAL1* and *TIA1* in *TIAL1*ko cells in the enriched Perturb-seq data. g) Gene expression of *TIAL1* and *TIA1* in *TIA1*ko cells in the enriched Perturb-seq data. h) Gene expression of *HSP90AA1* and *HSP90AB1* in *HSP90AA1*ko cells in the enriched Perturb-seq data. i) Gene expression of *HSP90AA1* and *HSP90AB1* in *HSP90AB1*ko cells in the enriched Perturb-seq data. Wilcoxon rank-sum P-values calculated and adjusted following differential gene expression analysis

As expected, most gene targets demonstrated reduced expression when targeted with sgRNAs **(Fig. 2b)**. A subset of paralog pairs exhibited subsequent upregulation of the intact or “response” paralog following loss of the target paralog **(Fig. 2b)**, suggestive of possible TA. Differential gene expression analysis confirmed the significant upregulation of response paralogs in six paralog pairs (Wilcoxon Rank Sum test, adjusted p-value <0.05) **(Fig. 2c)**.

Interestingly, this transcriptional adaptation was detected only in one direction; only loss of *NONO* induced *PSPC1* expression, not the reverse **(Fig. 2de)**, also observed in the whole transcriptome analysis **(Fig. 1g)**. This unidirectional TA was observed for all six paralog pairs that demonstrated TA **(Fig. 2d-i, S1h-m)**. sgRNA-level data showed varied activity dependent on sgRNA **(Fig. S2a)**, with most predicted to induce nonsense-mediated decay (NMD) based on known decay rules^21,22^ **(Fig S2bc**).

To determine if differences in paralog pair expression was an important factor in TA response, we evaluated the expression pattern of target paralogs in PC9 cells. Paralog pair expression was positively correlated (Pearson r=0.3, p-value=0.02) **(Fig S2d)** and the six pairs undergoing TA in our model did not demonstrate outlier expression, suggesting that large differences in paralog expression are not a driving force in transcriptional adaptation. These findings are among the first to present an unbiased screen of transcriptional adaptation among essential paralog pairs in human cells, identifying six independent paralog pairs that are susceptible to transcriptional adaptation following loss of one gene.

### Non-degradative CRISPR-mediated loss can induce transcriptional adaptation

Double-stranded breaks generated by Cas9 cutting induce frameshift mutations through insertions and deletions generated during DNA repair **(Fig. 3a)**. Frameshift mutations most often generate mRNA that will be degraded through NMD due to premature termination codons (PTCs)^21,22^. In contrast, CRISPR interference (CRISPRi) uses sgRNAs to localize transcriptional repression machinery to the transcription start site of target genes - silencing their expression, without generating mutant mRNA **(Fig. 3a)**. To assess whether the TA identified between the six paralog pairs observed in our CRISPRko dataset was unique to degradative loss, we evaluated the pan-genome CRISPRi screen performed in K562 cells by Replogle *et al*^23^. Interestingly, the majority of pairs did not show upregulation following CRISPRi-mediated loss of target paralogs **(Fig. 3b)**, despite robust cell and UMI coverage **(Fig. S3ab)**. Thus, these gene pairs are candidates for paralogs that undergo degradation-dependent TA, requiring the generation of mutant mRNA. However, two paralog pairs, *HSP90AA1-HSP90AB1* and *NONO-PSPC1*, exhibited upregulation in both the CRISPRko **(Fig. 1e)** and CRISPRi **(Fig. 3b,e)** experiments, indicating non-degradative mechanisms likely regulate adaptation of these pairs. As seen in the CRISPRko dataset **(Fig. 2f)**, a second *NONO* paralog, *SFPQ*, was also upregulated in response to CRISPRi mediated *NONO*-loss **(Fig. 3e)**. Further, paralog gene expression was highly correlated in K562 cells, again confirming that paralog gene expression patterns do not seem to predict which pairs undergo TA **(Fig S3c)**. These findings suggest that most of the TA observed in the CRISPRko dataset was likely degradation-dependent, with the exception of *HSP90AA1-HSP90AB1* and *NONO-PSPC1-SFPQ*.

**Figure 3:**
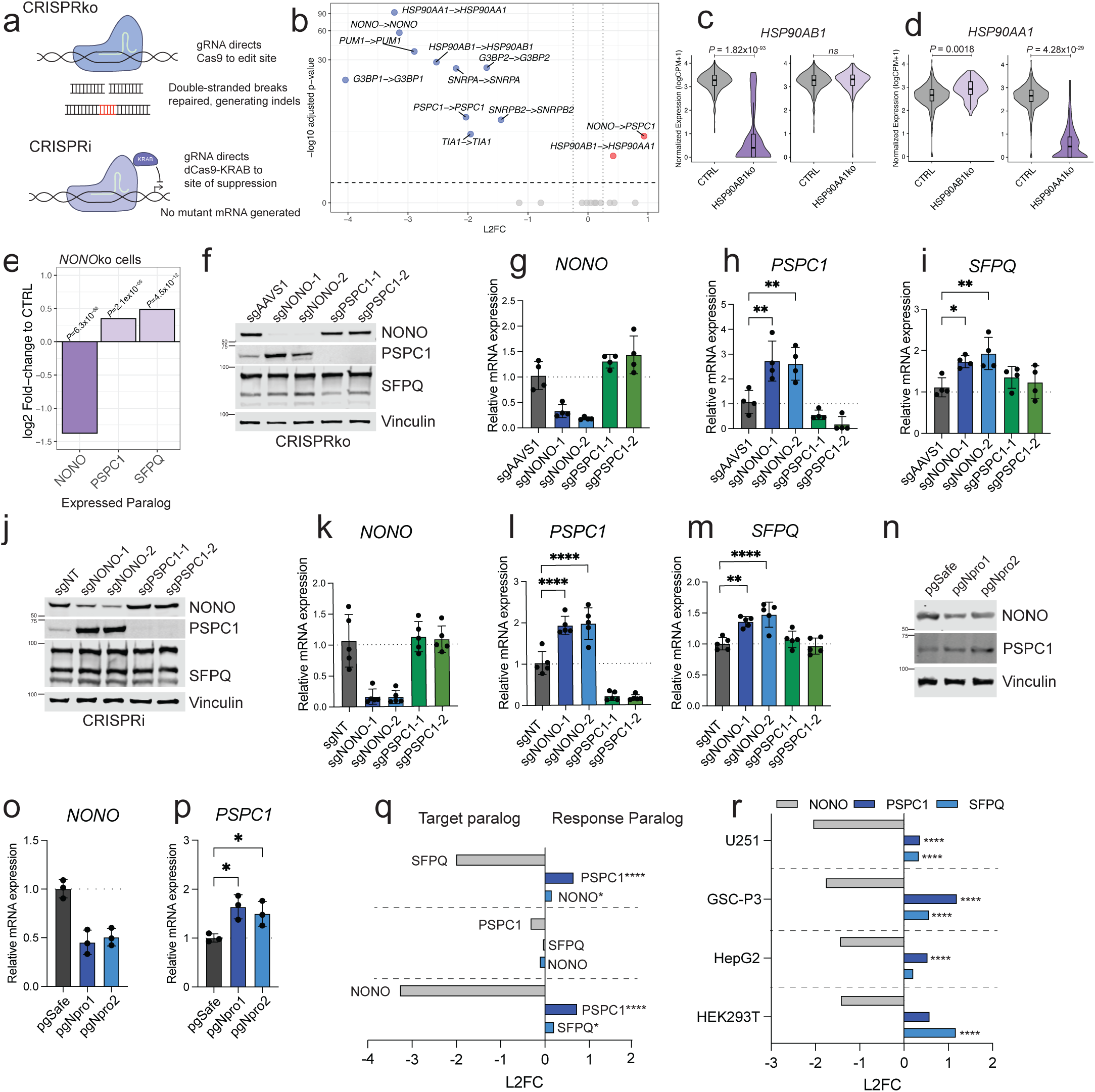
Paralog pair *NONO-PSPC1* undergo non-degradative transcriptional adaptation a) Schematic of CRISPRko versus CRISPRi gene silencing techniques. b) Selected target and response gene expression following pooled CRISPRi silencing of selected paralogs from the Replogle *et al* dataset, presented as “t*arget->response*” gene. A dashed line indicates adjusted p-value <0.05. c) Gene expression of *HSP90AA1* in Replogle *et al* Perturb-seq data by target gene knockout, Wilcoxon rank-sum adjusted P-values following differential gene expression analysis d) Gene expression of *HSP90AB1* in Replogle *et al* Perturb-seq data by target gene knockout, Wilcoxon rank-sum adjusted P-values following differential gene expression analysis e) Gene expression of *NONO*, *PSPC1*, *SFPQ* in *NONO*ko cells from Replogle *et al* CRISPRi screen, compared to cells containing control sgRNAs. Wilcoxon rank-sum adjusted P-values following differential gene expression analysis f) Western blot of NONO, PSPC1, and SFPQ following treatment with CRISPRko sgRNAs targeting safe locus *AAVS1*, two independent sgRNAs against *NONO*, and two independent sgRNAs against *PSPC1*. Representative of four independent replicates g) *NONO* mRNA expression by target CRISPRko sgRNA, 4 independent replicates. h) *PSPC1* mRNA expression by target CRISPRko sgRNA, 4 independent replicates, one-way ANOVA with Sidak’s multiple comparison test. **p<0.01. i) *SFPQ* mRNA expression by CRISPRko target sgRNA, 4 independent replicates, one-way ANOVA with Sidaks multiple comparison test, *p<0.05, **p<0.01 j) Western blot of NONO, PSPC1, and SFPQ following treatment with CRISPRi sgRNAs, control non-targeting and two independent sgRNAs against *NONO*, and two independent sgRNAs against *PSPC1* in PC9 cells expressing KRAB-ZIM3-dCas9, representative of four independent replicates. k) *NONO* mRNA expression by target CRISPRi sgRNA, 4 independent replicates. l) *PSPC1* mRNA expression by target CRISPRi sgRNA, 4 independent replicates, one-way ANOVA with Sidaks multiple comparison test, ****p<0.0001 m) *SFPQ* mRNA expression by target CRISPRi sgRNA, 4 independent replicates, one-way ANOVA with Sidaks multiple comparison test, ***p<0.001 ****p<0.0001 n) Western blot of NONO and PSPC1 following treatment with pgRNAs targeting *NONO* promoter in PC9 cells, representative of three independent replicates o) *NONO* mRNA expression following treatment with pgRNAs targeting NONO promoter in PC9 cells, three independent replicates. p) *PSPC1* mRNA expression following treatment with pgRNAs targeting NONO promoter in PC9 cells, three independent replicates, one-way ANOVA with Sidak’s multiple comparisons test, *p<0.05. q) *NONO*, *PSPC1*, and *SFPQ* mRNA expression from RNAseq in cells with CRISPRko of *SFPQ*, *PSPC1*, and *NONO* with response gene upregulation, Wald’s test with multiple hypothesis correction, *p<0.05, ****p<0.0001 r) *NONO*, *PSPC1*, and *SFPQ* mRNA expression from RNAseq in cells treated with siRNA targeting *NONO*, Wald’s test with multiple hypothesis correction ****p<0.0001

### Transcriptional adaptation of *NONO-PSPC1* also occurs through non-degadative silencing

We next tested whether CRISPRko and CRISPRi-mediated silencing of *NONO* can drive transcriptional responses in PC9 cells. Using CRISPRko, robust reduction of *NONO* **(Fig. 3fg)** induced upregulation of PSPC1 both at the protein **(Fig. 3f)** and mRNA (**Fig. 3h)** level, verifying TA following mutant mRNA induction seen in the Perturb-seq experiment **(Fig. 1ef and Fig. 2d)**. SFPQ followed a similar pattern **(Fig. 3f,i)**, with transcriptional upregulation evident in *NONO* knockout cells but not *PSPC1* knockout cells **(Fig. 3i)**. Using KRAB-ZIM3-dCas9^24^ **(Fig. 3a)** targeting both *NONO* and *PSPC1* through transcriptional silencing instead of genome editing, a strong induction of PSPC1 expression in *NONO*-inhibited cells was also observed at the protein **(Fig. 3j)** and mRNA **(Fig. 3kl)** level, validating the degradation-independent loss observed in the K562 CRISPRi screen^23^. Additionally, upregulation of *SFPQ* was observed after *NONO*-silencing at the mRNA level **(Fig 3jm)**. To further verify that mutant mRNA was not required for *NONO-PSPC1* TA, we generated promoter knockouts of *NONO*, which should prevent mRNA generation and thus prevent mRNA degradation. Indeed, promoter-loss of *NONO* using dual-targeting paired guide RNAs (pgRNAs) **(Fig. 3no, S3d)** induced upregulation of both PSPC1 protein **(Fig. 3n)** and mRNA **(Fig. 3p)**. These results demonstrate the strong paralog buffering within the *NONO-PSPC1-SFPQ* family, regardless of degradative or non-degradative gene expression silencing.

### *NONO-PSPC1-SFPQ* transcriptional adaptation is a broadly occurring phenomenon

Considering the robust upregulation of *PSPC1* and *SFPQ* in response to *NONO* loss, independent of how *NONO* loss was incurred, we next examined publicly available datasets that leverage both CRISPRko and siRNA downregulation of target genes. We assessed several published RNAseq datasets to determine whether this paralog adaptation was broadly occurring. In prostate cancer cell line 22Rv1^25^, CRISPRko loss of *NONO* and *SFPQ* induced upregulation of intact paralogs, while *PSPC1* loss did not **(Fig. 3q)**. Further, siRNA-mediated silencing of *NONO* across a panel of cancer cell lines U251^26^ (glioblastoma), GSC-P3^26^ (glioblastoma), HepG2^27^ (liver), and HEK293T^27^ all demonstrated upregulation of both *PSPC1* and *SFPQ*, further validating the robust and broadly occurring compensation among these paralogs **(Fig. 3r)**.

Thus, across at least 7 different cell lines using siRNA, CRISPRko and CRISPRi-mediated gene silencing methods, upregulation of *PSPC1* and often *SFPQ* is commonly observed after *NONO* inactivation. This tight regulation of the *NONO-PSPC1-SFPQ* family may be related to balancing their known essential cell functions in paraspeckles^28^, and more recently in recruiting telomerase to telomeres^29^. Recent work has also identified novel compounds that specifically target NONO using a unique cysteine residue not shared among the paralog family; interestingly, even pharmacological inhibition led to upregulation of both *PSPC1* and *SFPQ*^25^, further confirming that compensation in this paralog family can be induced without mutant mRNA.

### TIA1-TIAL1 TA is degradation-dependent in multiple cell line models

In contrast to *NONO-PSPC1*, we identified four paralog pairs that did not induce a measurable TA response in the CRISPRi system **(Fig. 3b)**, *TIA1-TIAL1*, *G3BP1-G3BP2*, *PUM1-PUM2*, and *SNRPA2-SNRPB* and hence may exhibit degradation-dependent TA. We selected *TIA1-TIAL1* to further investigate, due to the strong phenotype in the CRISPRko conditions **(Fig. 2fg)** and the classification of this gene pair as pan-synthetic lethal^30^, including in PC9 cells^4^. CRISPR knockout of *TIAL1* **(Fig. 4ab, S4a)** induced an upregulation of TIA1 at both the protein **(Fig. 4ab)** and mRNA **(Fig. 4c)** level, confirming our Perturb-seq findings. Conversely, CRISPRi-mediated, or non-degradative, loss of *TIAL1* did not significantly impact TIA1 protein expression **(Fig. 4ab)**, or mRNA expression **(Fig. 4c)** despite robust *TIAL1* gene silencing. To determine whether these patterns were cell-type specific or shared across cell contexts, we repeated the assays in another lung adenocarcinoma cell line, Calu6. As in PC9 cells, degradation-dependent upregulation of TIA1 following *TIAL1* loss was also observed **(Fig. 4d-f, S4b)**.

**Figure 4:**
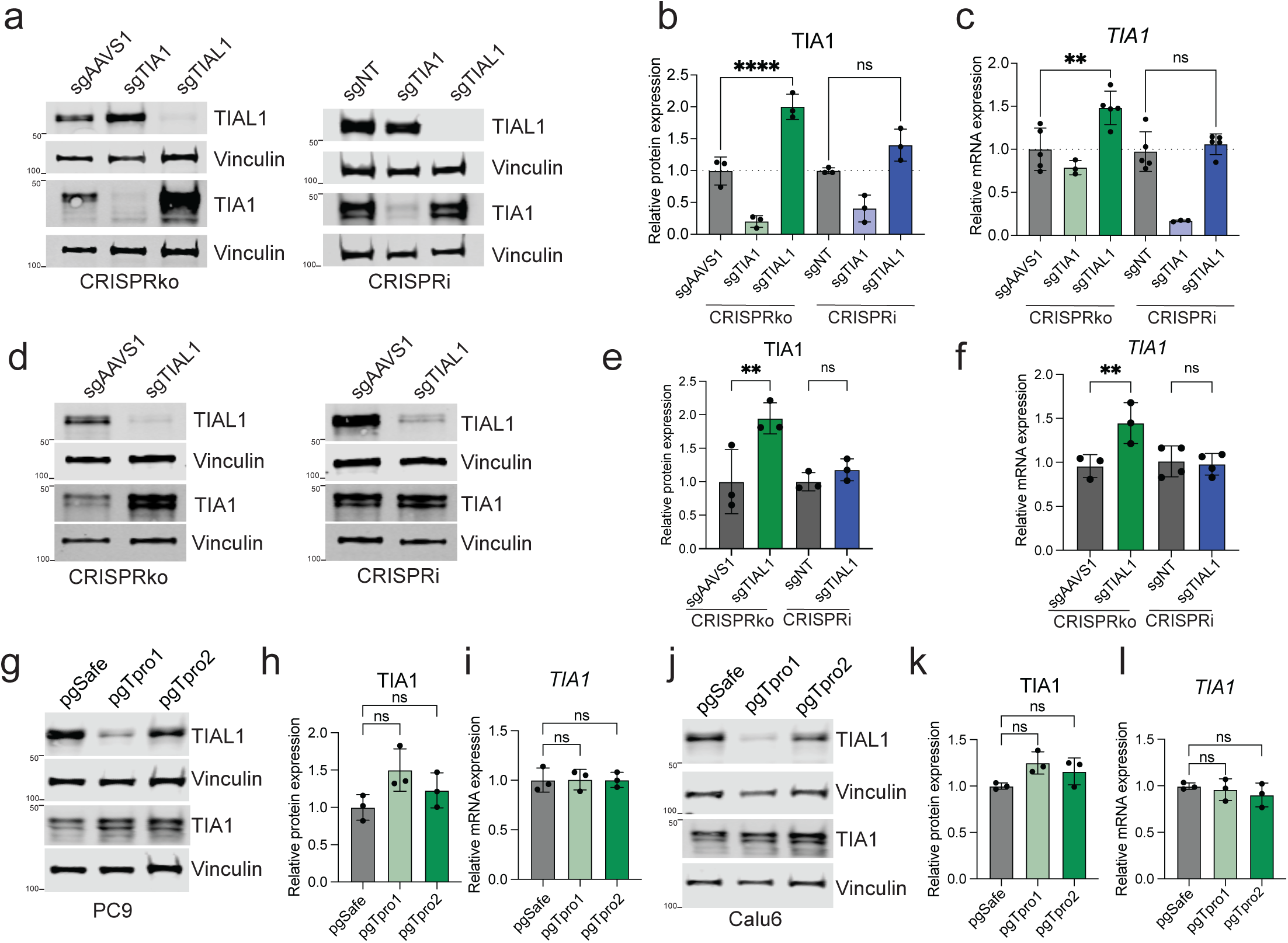
T*I*A1*-TIAL1* transcriptional adaptation is degradation-dependent a) Western blot of TIA1 and TIAL1 in PC9 cells following treatment with CRISPRko (right) or CRISPRi (left) sgRNAs, representative of three biological replicates. b) Quantification of western blots in a, one-way ANOVA with Sidak’s multiple comparisons test, ****p<0.0001 c) *TIA1* mRNA expression in PC9 cells following treatment with indicated sgRNAs, one-way ANOVA with Sidak’s multiple comparisons test, **p<0.01 d) Western blot of TIA1 and TIAL1 in Calu6 cells following treatment with CRISPRko or CRISPRi sgRNAs, representative of three independent replicates. e) Quantification of Calu6 western blots in d, one-way ANOVA with Sidak’s multiple comparisons test, **p<0.01. f) *TIA1* mRNA expression in Calu6 cells following treatment with indicated sgRNAs, three independent replicates, one-way ANOVA with Sidak’s multiple comparisons test, **p<0.01 g) Western blot of TIA1 and TIAL1 in PC9 cells following knockout of *TIAL1* promoter using pgRNAs, representative of three independent replicates h) Quantification of TIA1 from western blot in g, three independent replications, one-way ANOVA with Sidak’s multiple comparison test. i) *TIA1* mRNA expression in PC9 cells with TIAL1 promoter knockout using pgRNAs, three independent replicates, one-way ANOVA with Sidak’s multiple comparisons test. j) Western blot of TIA1 and TIAL1 in Calu6 cells following knockout of *TIAL1* promoter using pgRNAs, representative of three independent replicates k) Quantification of TIA1 protein expression from western blot in j, three independent replicates, one-way ANOVA with Sidak’s multiple comparison test. l) *TIA1* mRNA expression in Calu6 cells with TIAL1 promoter knockout using pgRNAs, three independent replicates, one-way ANOVA with Sidak’s multiple comparisons test.

To further validate the dependence of TIA1 upregulation on the degradative-loss of *TIAL1*, we generated pgRNAs targeting the promoter of *TIAL1,* which reduced TIAL1 expression without generating mutant mRNA **(Fig. S4c)**. pgRNAs demonstrated varying degrees of knockout efficiency **(Fig. 4i, S4de)**, and while a small and non-significant upregulation of TIA1 protein levels was observed **(Fig. 4ij)**, no upregulation of *TIA1* mRNA followed promoter loss of *TIAL1* in both PC9 cells **(Fig. 4k)** and Calu6 cells **(Fig 4l-n, S4fg)**. Together, these data show that upregulation of TIA1 in response to CRISPRko-mediated loss of its paralog *TIAL1* occurs via a degradation-dependent mechanism.

### *TIA1-TIAL1* TA requires nonsense-mediated decay

To further assess whether the degradation-dependent TA between *TIA1-TIAL1* requires the NMD pathway, we determined whether blocking NMD could prevent TA of *TIA1* following *TIAL1* knockout. SMG1 is the first kinase activated in response to ribosome stalling on PTCs^31^. SMG1 directly phosphorylates UPF1, which is required for the subsequent recruitment of nucleases, including SMG6^32^, that degrades PTC-bearing mRNA **(Fig 5a)**. Treatment with SMG1 inhibitor (hSMG1i-11j) for 8 hours reduced phosphorylation of its target, UPF1 at Ser1197 **(Fig. 5b)**, as expected, with minimal effect on total UPF1 protein levels. mRNA expression of *TIA1* was restored to baseline levels following SMG1i treatment, suggesting that blocking NMD prevents TA between *TIA1-TIAL1* **(Fig. 5c)**. Further, a significant restoration of *TIAL1* mRNA was observed **(Fig. 5d)**, suggesting that by blocking NMD, degradation of mutant *TIAL1* mRNA was also blocked. A similar effect was observed in Calu6 cells **(Fig. 5e)**, where treatment with SMG1i markedly reduced phospho-UPF1 and blocked the transcriptional response of *TIA1* in response to *TIAL1* mutation **(Fig. 5fg)**. Next, to assess whether core components of the NMD pathway are genetically required for this phenomenon, we treated both control and *TIAL1*ko cells with a combination of siRNAs targeting both UPF1 and SMG6, previously identified to mute TA responses in a model of Duchenne’s muscular dystrophy^14^ and required for the recruitment of RNA degradation nucleases, including SMG6 **(Fig. 5a)**.

**Figure 5:**
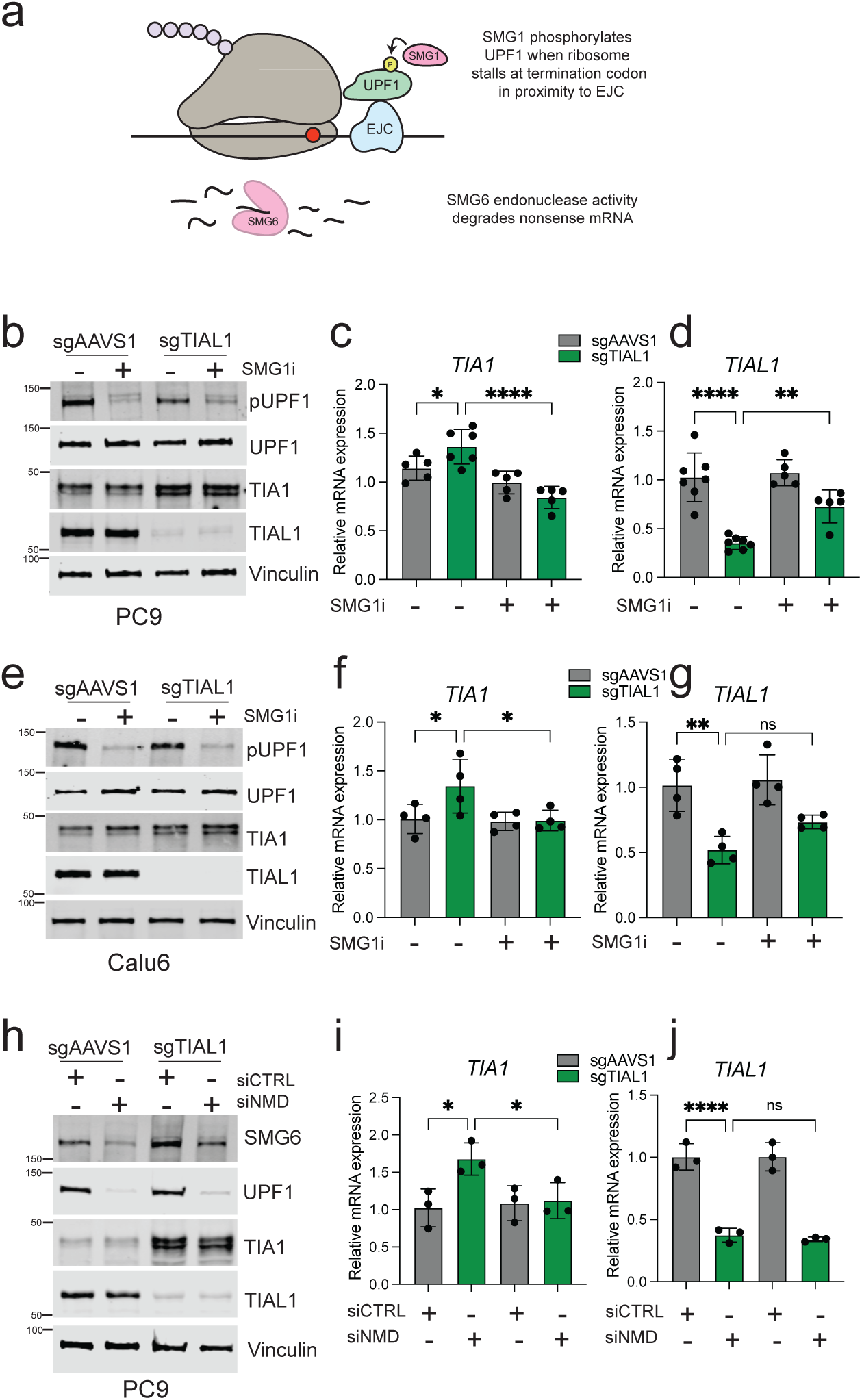
TIA1-TIAL1 degradation-dependent TA requires NMD a) Schematic representation of nonsense-mediated decay following detection of a premature termination codon. EJC = exon junction complex. b) Western blot of lysates following treatment of indicated PC9 cells with 500nM SMG1i for 8h, representative of three independent replicates. c) *TIA1* mRNA expression following treatment of indicated PC9 cells with 500nM SMG1i for 8h, five independent replicates, one-way ANOVA with Sidak’s multiple comparison test, *p<0.05, ****p<0.0001. d) *TIAL1* mRNA expression following treatment of indicated PC9 cells with 500nM SMG1i for 8h, five independent replicates, one-way ANOVA with Sidak’s multiple comparison test, **p<0.01, ****p<0.0001. e) Western blot of lysates following treatment of indicated Calu6 cells with 500nM SMG1i for 8h, representative of three independent replicates. f) *TIA1* mRNA expression following treatment of indicated Calu6 cells with 500nM SMG1i for 8h, four independent replicates, one-way ANOVA with Sidak’s multiple comparison test, *p<0.05. g) *TIAL1* mRNA expression following treatment of indicated Calu6 cells with 500nM SMG1i for 8h, four independent replicates, one-way ANOVA with Sidak’s multiple comparison test, **p<0.01. h) Western blot of PC9 cells treated with siRNAs targeting SMG6 and UPF1, harvested 72h post-transfection. Representative of three independent replicates i) *TIA1* mRNA expression following treatment with indicated siRNAs for 72h, three independent replicates, one-way ANOVA with Sidak’s multiple comparison test, *p<0.05 j) *TIAL1* mRNA expression following treatment with indicated siRNAs for 72h, three independent replicates, one-way ANOVA with Sidak’s multiple comparison test, ****p<0.001

Reduction of both UPF1 and SMG6 **(Fig. 5h, S4hi)** had a minimal effect on response gene TIA1 protein levels **(Fig. 5h)**, but reduced *TIA1* mRNA levels back to control levels **(Fig. 5i)**, without an appreciable impact on *TIAL1* mRNA levels **(Fig. 5j)**. We confirmed that activity of the ribosome was important in this degradation-dependent phenotype; treatment with the ribosome inhibitor cycloheximide blocked *TIA1* upregulation, and stabilized mutant *TIAL1* mRNA **(Fig. S4jk)**. This work demonstrates that both pharmacological and genetic inhibition of NMD blocks the TA response between *TIAL1* and *TIA1* paralogs, further confirming that the TA between these genes is indeed degradation-dependent.

### Degradation-dependent TA remodels chromatin of the response gene

Previous reports have suggested that the mechanism of degradation-dependent TA involves nuclear re-entry of degraded mRNA fragments to upregulate related genes in a sequence-driven manner^11^. To understand the role of chromatin remodelling in the degradation-dependent TA occurring between *TIA1-TIAL1*, we first assessed whether chromatin at the *TIA1* promoter changed in response to *TIAL1*ko. Using chromatin immunoprecipitation paired with qPCR, we identified that H3K4me3, a histone modification associated with active transcription, was significantly enriched at the *TIA1* promoter following degradative loss of *TIAL1* **(Fig. 6a)**, while control gene occupancy was unaffected **(Fig. S4l)**. These results confirm that active transcription is a key component of the TA response and suggest that the upregulation of adapting genes may be occurring through chromatin remodelling at the response gene promoters.

**Figure 6:**
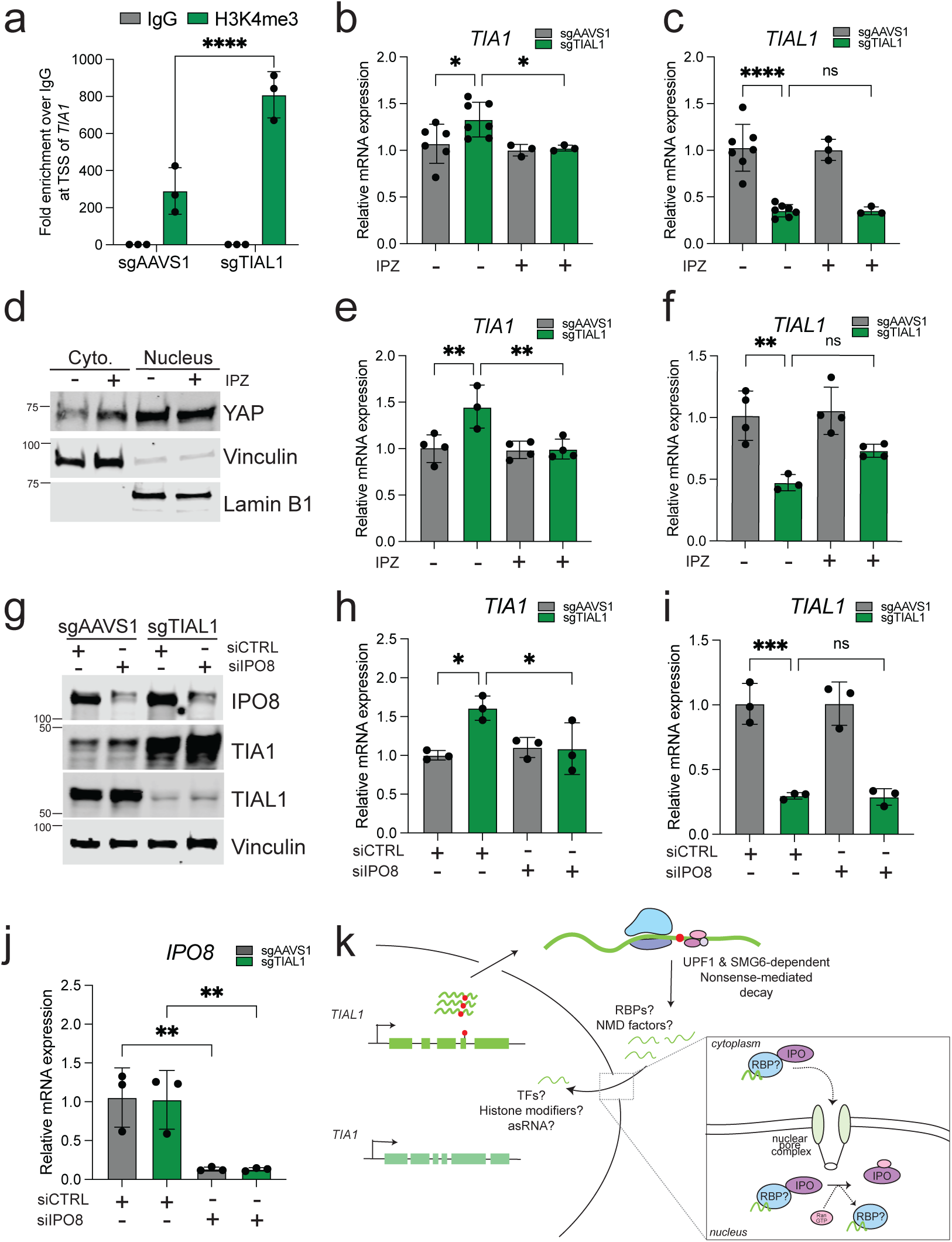
Degradation-dependent TA requires active nuclear import a) Chip-PCR at promoter of TIA1 of fold enrichment of activating histone mark H3K4me3 compared to IgG following treatment with control sgAAVS1 or sgTIAL1 sgRNAs in PC9 cells. Three independent replicates, two-way ANOVA, ****p<0.0001 b) *TIA1* mRNA expression in PC9 cells following treatment with IPZ at 20μM for 8h, 3-5 independent replicates, one-way ANOVA with Sidak’s multiple comparisons test, *p<0.05. c) *TIAL1* mRNA expression in PC9 cells following treatment with IPZ at 20μM for 8h, 3-5 independent replicates, one-way ANOVA with Sidak’s multiple comparisons test, ****p<0.0001. d) Western blot of nuclear and cytoplasmic lysates from PC9 cells treated with 20μM IPZ for 8 hours. e) *TIA1* mRNA expression in Calu6 cells following treatment with IPZ at 20μM for 8h, 3-5 independent replicates, one-way ANOVA with Sidak’s multiple comparisons test, **p<0.01. f) *TIAL1* mRNA expression in Calu6 cells following treatment with IPZ at 20μM for 8h, 3-5 independent replicates, one-way ANOVA with Sidak’s multiple comparisons test, **p<0.01. g) Western blot of PC9 cells treated indicated sgRNAs and with indicated siRNAs for 72h. Representative of three independent replicates. h) *TIA1* mRNA expression in PC9 cells following treatment with indicated siRNAs for 72h, three independent replicates, one-way ANOVA with Sidak’s multiple comparisons test, *p<0.05 i) *TIAL1* mRNA expression in PC9 cells following treatment with indicated siRNAs for 72h, three independent replicates, one-way ANOVA with Sidak’s multiple comparisons test, ***p<0.001 j) *IPO8* mRNA expression in PC9 cells following treatment with indicated siRNAs for 72h, three independent replicates, one-way ANOVA with Sidak’s multiple comparisons test, **p<0.01 k) Schematic of degradation-dependent transcriptional adaptation occurring between *TIAL1-TIA1* paralogs. NMD = nonsense mediated decay, RBP = RNA binding protein, TF = transcription factor, asRNA = antisense RNA, IPO = importin.

### TIA1-TIAL1 degradation-dependent TA requires active nuclear import

Despite current mechanistic models of TA alluding to the requirement of re-entry of degraded mRNA fragments to the nucleus, to date, none have assessed whether this requires active nuclear import. To address this, we treated control and *TIAL1*ko cells with the importin-ꞵ1 inhibitor, importazole^33^ (IPZ). Indeed, the activity of importin-ꞵ1s was required for the degradation-dependent TA response; following inhibition of nuclear import, the *TIA1* mRNA upregulation induced by *TIAL1*ko response was returned to control levels in both PC9 **(Fig. 6bc)**. Assessment of the nuclear-import responsive protein YAP confirmed suppression of nuclear import under these treatment conditions **(Fig 6d)**. Calu6 cells demonstrated a similar response **(Fig. 6ef)**.

We next hypothesized that known RNA silencing components could be involved in the TA response between *TIA1-TIAL1*. Given the role of argonaute proteins in binding RNA and regulating mRNA abundance, as well as previously reported roles of AGO2 specifically altering transcriptional landscapes^34–36^, we assessed whether loss of AGO2 impacts the TA response. However, *AGO2* suppression by siRNAs did not significantly affect the degradation-dependent TA response between *TIA1-TIAL1*, despite a robust reduction in *AGO2* mRNA expression **(Fig. S4m-o)**. In contrast, silencing of *IPO8*, one of the previously reported nuclear importins of AGO2^37,38^, did reduce the degradation-dependent TA response to control levels **(Fig. 6g-j)**.

Thus, while AGO2 does not appear to be the relevant importin cargo, active nuclear import through importing proteins was required for degradation-dependent TA **(Fig. 6k)**.

These pharmacological and genetic disruption studies show that active nuclear import is an essential component of degradation-dependent TA, without affecting mutant mRNA degradation. Further work is required to identify the proteins responsible for degraded mRNA transport into the nucleus and how chromatin remodelling occurs to activate the target gene.

## Discussion

Degradation-dependent transcriptional adaptation is a newly uncovered gene regulation program that appears to contribute to cell fitness via genetic compensation. Despite the broad consequences of this regulatory mechanism on gene expression across disease types, and the utility of the CRISPRko-based therapeutics, little work has been done to assess how broadly this phenomenon occurs, particularly among paralog families.

Here, we leveraged Perturb-seq to assess the frequency of TA among highly essential paralog genes in human cells. Surprisingly, we find that TA is relatively selective; despite high levels of homology between gene pairs, TA measurable within our system occurred in six of 54 (∼11%) pairs assessed **(Fig. 2c, 2d-i, S1h-m)**. Since little is known about the dynamics of adaptation, it is possible that the selected timepoint to assess TA responses affected our ability to detect some TA phenotypes. We chose a long treatment with sgRNAs to verify robust knockout of target genes **(Fig. S1a)**, which could mask initial TA responses that occur early.

Cell coverage in Perturb-seq **(Fig S1bc)** may also limit the detection of more muted TA responses, though our use of hybrid-capture enrichment for target genes partially mitigates this concern **(Fig. S1g)**.

Exploring the mechanism of TA of paralogs, we assessed both non-degradative and degradation-dependent models of TA. Two of the pairs we identified to undergo TA, *NONO-PSPC1* and *HSP90AA1-HSP90AB1*, also exhibited TA in a non-degradative assay **(Fig. 3b-e)**. These findings demonstrate that though CRISPRi-based gene silencing tools may bypass degradation-dependent TA, non-degradative TA is also occurring among paralog genes and likely contributes to buffering commonly observed in genetic studies of paralogs.

Extended characterization of degradation-dependent TA among RNA-stress granule associated paralogs *TIA1-TIAL1*^39^ demonstrated a requirement for NMD, through both pharmacological **(Fig. 5b-c)** and genetic inhibition **(Fig. 5h-j)** of NMD. For the first time, we report that active nuclear import through importin-β1 is required for this phenomenon **(Fig. 6b-f)**, making it clear that re-entry of degraded fragments into the nucleus to upregulate sequence-specific response genes is occurring. In our model, importin 8 seems to be responsible for this import **(Fig. 6g-j)**, but further testing of other importin proteins in other models is important to assess whether importin 8’s function is cell line or context-dependent **(Fig. 6k)**.

Very recent work uncovered an RNA-binding protein (RBP), ILF3, as necessary for TA responses in both mouse and human model systems^12^. Whether the role of this RBP is shared across multiple cell contexts is not clear, and assessing the function of ILF3 and its functional cooperation with importin-β1 in the regulation of paralog TA is an important future direction.

Overall, this work sheds light on transcriptional adaptation among highly essential paralog genes in human cells and provides the first systematic assessment of transcriptional adaptation among paralog genes. This work also adds mechanistic insight into this newly discovered gene regulatory mechanism and its dependence on active nuclear import.

## Methods

### Cell culture & treatments

PC9, Calu6 and HEK293T were all obtained from the ATCC. PC9 cells were grown in RPMI (Gibco) + 10% FBS, Calu6 cells were grown in EMEM (ATCC) + 10% FBS, and HEK293T cells were grown in DMEM (Genesee Scientific) + 10% FBS, all at 37°C and 5% CO2. Cell lines were routinely tested for mycoplasma and validated via STR profiling at the Fred Hutch Genomics core. Cells were treated with the following concentrations of small molecule: cycloheximide (Cell Signalling Technologies, #2112) at 100mg/mL for 8 hours. hSMG1-11j (MedChemExpress, HY-124719) at 500nM for 8 hours. Importazole (MedChemExpress, HY-101091) at 20μM for 8 hours. KRAB-ZIM3-dCas9 (Addgene #154472) PC9 and Calu6 cells were generated by infecting cells with lentivirus in the presence of 8 μg/ml polybrene (Sigma Aldrich, TR-1003).

Following infection, cells were selected for 3-10 days in 10μg/mL blasticidin (Gibco, A1113903).

### Lentivirus preparation & selection

HEK293T cells were transfected with 6μg pSPAX (Addgene #12260), 2μg pMD.2G (Addgene #12259) and 6.5μg of vector, using 46μg of PEI-MAX (Polysciences, 24765-100). CRISPRko gRNAs and pgRNAs were cloned into LentiCrispr_V2 (Addgene #52961), CRISPRi gRNAs were cloned into pLentiguide (Addgene #52963). pgRNAs were cloned as previously described^4^, geneblock synthesis performed by Azenta. Both PC9 and Calu6 cells were infected with lentivirus in the presence of 8μg/mL polybrene. Following infection, cells were selected in 2μg/mL puromycin (Gibco, A1113803) for 72h.

### sgPEN library design & cloning

Paralog pairs were nominated for inclusion based on genetic interaction scores from a previously published pgRNA screen in PC9 cells^4^. Oligos were synthesized at Twist, and cloned using standard procedures as described^40^ into pLentiGuide (Addgene #52963).

### sgPEN dropout screen

PC9-Cas9 cells were screened as previously described^42^, briefly, sgPEN lentivirus at an MOI of ∼0.3, in 8μg/mL polybrene. The following day, cells were selected in 2ug/mL puromycin for 72h. Following selection, cells were counted and split into three independent replicates. The screen was passed for 18 days, with cell pellets collected at each passage. gDNA was extracted from cell pellets at T0 and T25 using Qiagen DNeasy Blood and Tissue miniprep kit (Qiagen, 69504). sgRNAs were amplified from 1 μg of gDNA using KAPA HIFI master mix, using manufacturer-recommended cycling conditions. Sequencing libraries were sequenced on an Illumina Mi-Seq machine at 1000X read depth. sgRNA distribution was assessed using MAGeCK.

### sgPEN single cell preparation, library prep & sequencing

Perturb-seq samples were generated from T18 cells using the Chromium Next GEM Single Cell 5’ Reagent Kits v2 (Dual Index) with Feature Barcode technology for CRISPR screening (10x Genomics, 1000264, 1000267, 1000287) 5’ CRISPR kit (10X genomics, 100451), and the targeted gene expression kit (10X genomics, discontinued). ∼10,000 cells/ GEM well were targeted, with 12 total reactions. scRNA-seq transcriptome and gRNA libraries were prepared as per the manufacturer’s directions. ∼300ng of each transcriptome library was moved forward for Targeted Gene Expression, using a custom bait library generated by IDT, gRNA, whole transcriptome, and targeted gene expression libraries were sequenced on an Illumina NextSeq as per manufacturer’s instructions.

### Single-cell analysis

10x Genomics Cell Ranger count v7.1.0 was used to align scRNA-seq reads to the human (GRCh38) 2020-A reference, providing target enrichment panel sequences and feature reference library of sgRNAs. Reactions were aggregated using Cell Ranger Aggr v7.1.0. Further analysis was performed in Seurat v5. Cells were first filtered for single gRNA calls per cell, assigning sgRNA calls per cell if one sgRNA was >70% of total reads, for either the enriched or whole transcriptome samples. Next, the standard Seurat pipeline for normalization and clustering was performed^43^. Briefly, cells were excluded if they contained < 500 UMIs, detected < 500 genes, or had > 4% mitochondrial DNA reads. Cells were finally selected based on detection in both the enriched and whole-transcriptome datasets, with a final count of 36132 cells total. gRNA guide efficiency was first assessed by pseudobulking scRNAseq counts by sgRNA target gene and comparing to a control pseudobulk containing safe or NTC sgRNAs. To assess target and response gene expression, FindMarkers^43^ was run between all cell clusters based on the target gene, comparing to control cells with safe or NTC gRNAs.

### Sequence similarity analysis

Differential gene matrices were generated as described above, providing a table of target gene and response gene pairs, with corresponding LFC and Wilcoxon P-values. MANE-select mRNA sequences for all target genes were collected from Ensembl, capturing the 5’ UTR, coding sequence, and 3’ UTR. DNA sequences for all response genes was collected from Ensembl, capturing the entire annotated gene plus 2Kb upstream of the transcriptional start site, using pyfaidx^44^ to index the GRCh38 assembly. Pairwise mRNA-DNA alignment scores were calculated using the Biopython Bio.Align pairwise alignment using preset parameters for blast alignment scoring; alignment scores were collected for all 6 comparisons, split by genomic feature: 5’UTR, CDS, 3’UTR, predicted promoter, and DNA gene sequence.

Response gene hits were selected based on an L2FC > 0.5 and FDR < 0.1 from all target gene clusters. Control non-response genes were selected based on minimal effect, using cutoffs of L2FC<0.1 and L2FC-0.1, then sub-sampling 10,000 response genes randomly across all target gene clusters without replacement.

### Analysis of publicly available RNA-seq datasets

Raw sequencing data were retrieved from GEO datasets^25,27^ using a WILDS WDL module for RNA-seq transcript quantification. (https://github.com/getwilds/wilds-wdl-library/tree/1753e45fd0de71d759250c176f95dd9a9cd6e988/modules/ww-salmon). Read counts were generated using Salmon quasi-mapping. To calculate differential gene expression, pytximport was used to import Salmon quantification data and discretize counts, and pydeseq2 was used to generate different gene expression profiles.

### Western blotting & quantification

Lysates were collected using 4X Laemmli buffer (Bio-Rad, 1610747), homogenized through repeated passages through a 20g needle, and boiled at 95°C for 5 minutes. ∼15μg of protein was loaded per lane of BioRad precast 4-15% polyacrylamide gels (Bio-Rad, 4561084), and gels were run at 120V for ∼70 minutes in 1X running buffer (Bio-Rad, 1610732). Gels were transferred to ethanol-activated PVDF using the BioRad Turboblot system (Bio-Rad, 1704150). Membranes were blocked for 1h, shaking at room temperature using LiCOR PBS-T blocking buffer (Licor, 927-70003). Primary antibodies were incubated overnight at 4°C in blocking buffer. The following day, membranes were washed using 1X TBS-T (Genesee, 18-235B) and incubated in secondary antibody in blocking buffer for 1h at room temperature. Following 3-4 washes in TBS-T and a final rinse in PBS, blots were imaged on the LI-COR Odyssey imaging system (LI-COR Biosciences). Western blot quantification was performed by densitometry, using ImageJ^45^, and normalized to loading controls.

### Quantitative PCR

Lysates were collected in Trizol (Invitrogen, 10296010), and RNA was isolated using the Zymo RNA mini prep kit (Zymo, ZR2072), with on-column DNase digestion. 1-2 μg of RNA was used to generate cDNA using the High-Capacity cDNA Reverse Transcription kit (ThermoFisher, 4368814). cDNA was diluted and amplified with target-specific primers using the PowerUP SYBR green master mix (ThermoFisher, A25742) on a Biorad CFX-384 PCR machine (Bio-Rad, 1855484). Relative fold change was calculated using the delta-delta CT method, normalized to cyclophilin B.

### ChIP-qPCR

PC9 cells were infected and selected with corresponding gRNAs. Following selection, cells were subjected to ChIP-qPCR using the SimpleChIP Plus Sonication Chromatin IP kit (Cell Signalling Technologies, #56383). Briefly, cells were fixed in a final concentration of 1% formaldehyde (Fisher Scientific, RSOF0010500) for 8 minutes at room temperature, and quenched in 125mM glycine for 5 minutes at room temperature. Cells were collected by scraping, centrifuged, and stored at-80°C for further processing. Following cell permeabilization by the manufacturer’s protocol, lysates were sonicated at 30% amplitude, 30s on 30s off for 25 minutes on a VWR probe sonicator on ice (VWR, 76193-590). 10μg of sheared DNA was subjected to immunoprecipitation with the described antibodies. Immunoprecipitation was performed as per the manufacturer’s instructions, and DNA was eluted & quantified. Target sequences were amplified using PowerUP SYBR Master Mix, and target-specific primers (see Supp Table 4). All samples were normalized to IgG input.

### siRNA knockdowns

The following siRNAs were purchased from Horizon Discovery, UPF1 (Custom, CAGCGGAUCGUGUGAAGAA), SMG6 (Custom, CCAGUGAUACAGCGAAUUA), IPO8 (J-012256-19-0005), AGO2 (J-004639-06-0005) non-targeting pool (D-001810-10-05). siRNAs were transfected into PC9 cells using RNAiMAX (ThermoFisher, 13778030) in Opti-MEM (Gibco, 31985062), at a 15nM final siRNA concentration. 72h post-transfection, lysates were collected for western blotting and RNA extraction.

### Nuclear/cytoplasmic fractionations

Fractionation was performed using the NE-PER Nuclear and Cytoplasmic extraction kit (ThermoFisher, 78833) following the manufacturer’s instructions. Lysates were quantified using the Pierce BCA protein assay (ThermoFisher, 23227). Western blots were run as described.

## Acknowledgements

The authors would like to thank all members of the Berger lab for their support and helpful discussions, especially Elana Thieme and Saksham Gupta. We also thank Dr. Sitapriya Moorthi and Dr. Nevraj Keijou for discussions and critical feedback on the manuscript. SO is a Washington Research Foundation postdoctoral fellow. MHF is supported by the NIH award R25HG012337. This research was supported in part by NCI R01CA262556 and the American Cancer Society. This research was supported by NIH P30 CA015704 of the Fred Hutch/University of Washington/Seattle Children’s Cancer Consortium, which includes the Genomics & Bioinformatics Shared Resource

---

**Supplementary Figure 1.**
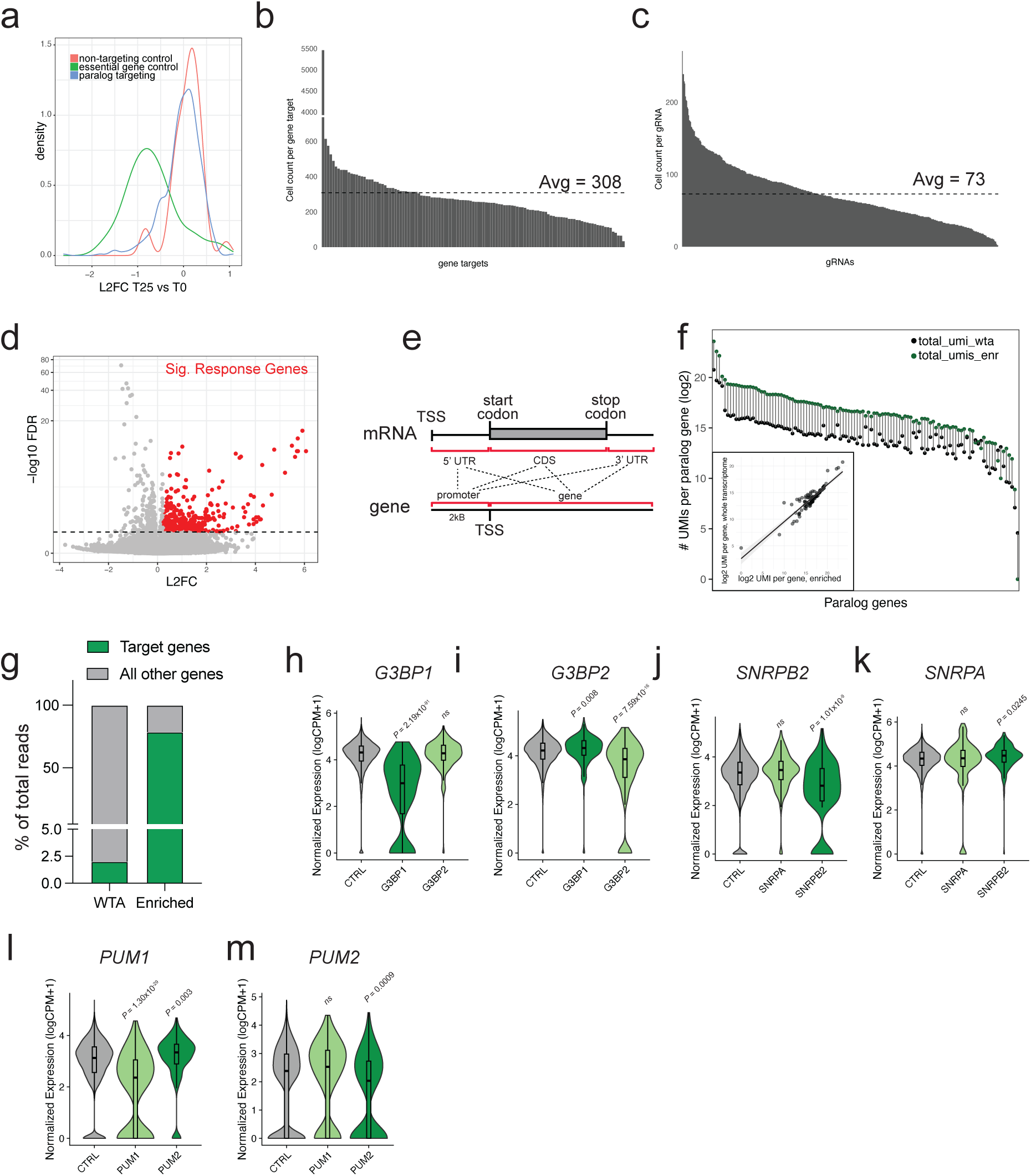
a) sgRNA distribution of T25 vs T0 of sgPEN test screen, labelled by sgRNA type. b) Cell coverage in Perturb-seq dataset by target gene. c) Cell coverage in Perturb-seq dataset by sgRNA. d) Differential gene expression results across all target knockout genes, significant response genes colored in red and selected for further study. e) Schematic of mRNA/DNA element sequence comparisons in Fig. 1h. f) UMI enrichment between hybrid capture and whole transcriptome datasets. Inset is correlation plot of UMI counts per gene between enriched and whole transcriptome dataset. g) Read enrichment of targeted genes in hybrid capture enrichment versus whole transcriptome read coverage. h) *G3BP1* mRNA expression in enriched Perturb-seq data, separated by target gene knockout. Wilcoxon rank-sum P-values calculated and adjusted following differential gene expression analysis i) *G3BP2* mRNA expression in enriched Perturb-seq data, separated by target gene knockout. Wilcoxon rank-sum P-values calculated and adjusted following differential gene expression analysis j) *SNRPB2* mRNA expression in enriched Perturb-seq data, separated by target gene knockout. Wilcoxon rank-sum P-values calculated and adjusted following differential gene expression analysis k) *SNRPA* mRNA expression in enriched Perturb-seq data, separated by target gene knockout. Wilcoxon rank-sum P-values calculated and adjusted following differential gene expression analysis l) *PUM1* mRNA expression in enriched Perturb-seq data, separated by target gene knockout. Wilcoxon rank-sum P-values calculated and adjusted following differential gene expression analysis m) *PUM2* mRNA expression in enriched Perturb-seq data, separated by target gene knockout. Wilcoxon rank-sum P-values calculated and adjusted following differential gene expression analysis

**Supplementary Figure 2.**
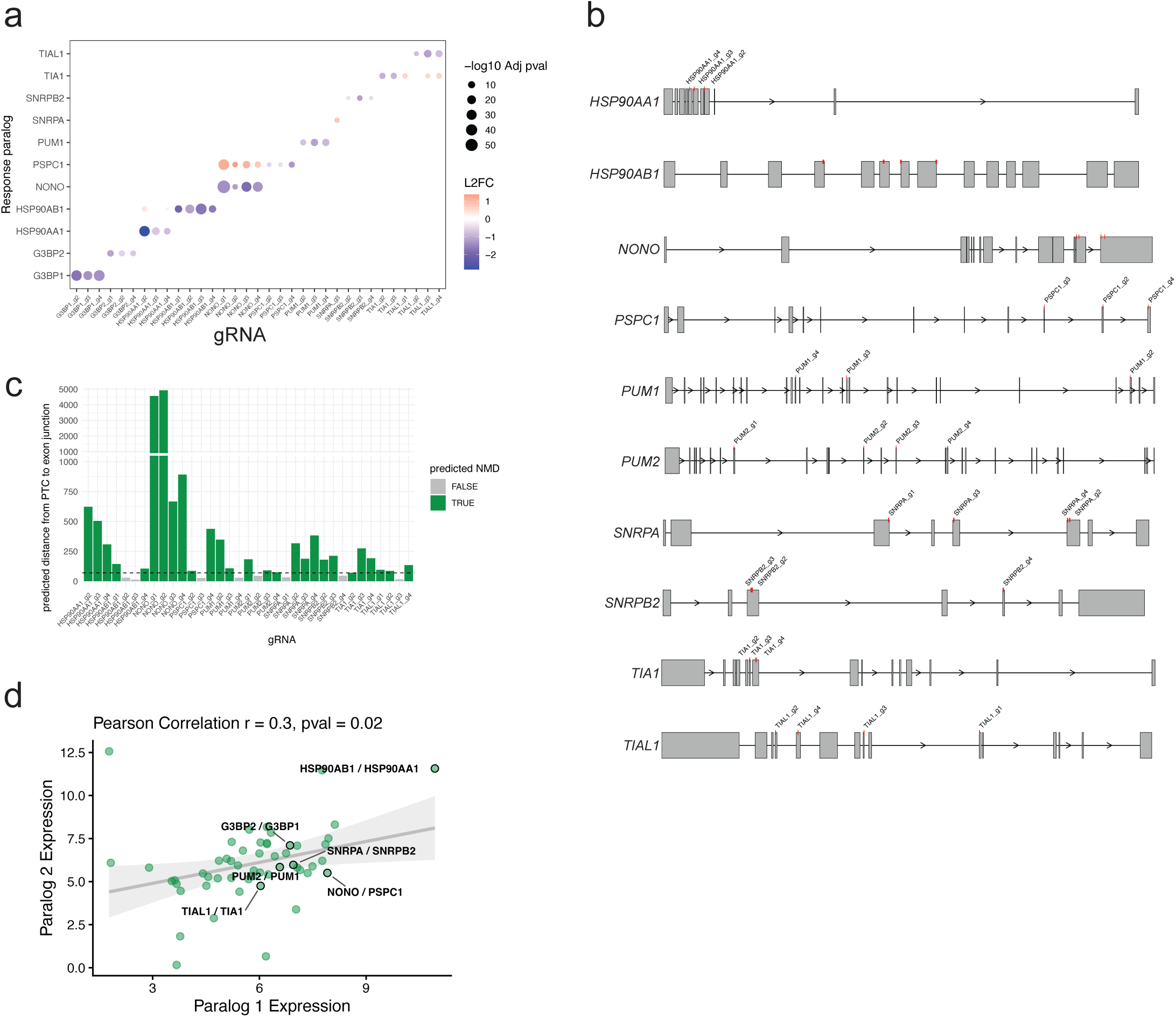
a) gRNA activity against target genes and corresponding response genes, enriched dataset. Wilcoxon rank sum P-values, adjusted by multiple hypothesis testing. b) Schematic of gRNA target positions across all genes detected to undergo transcriptional adaptation. c) Predicted distance from CRISPR cut site to the nearest exon junction for each gRNA targeting paralogs that undergo TA. d) Correlation of paralog pair expression in control PC9 cells.

**Supplementary Figure 3.**
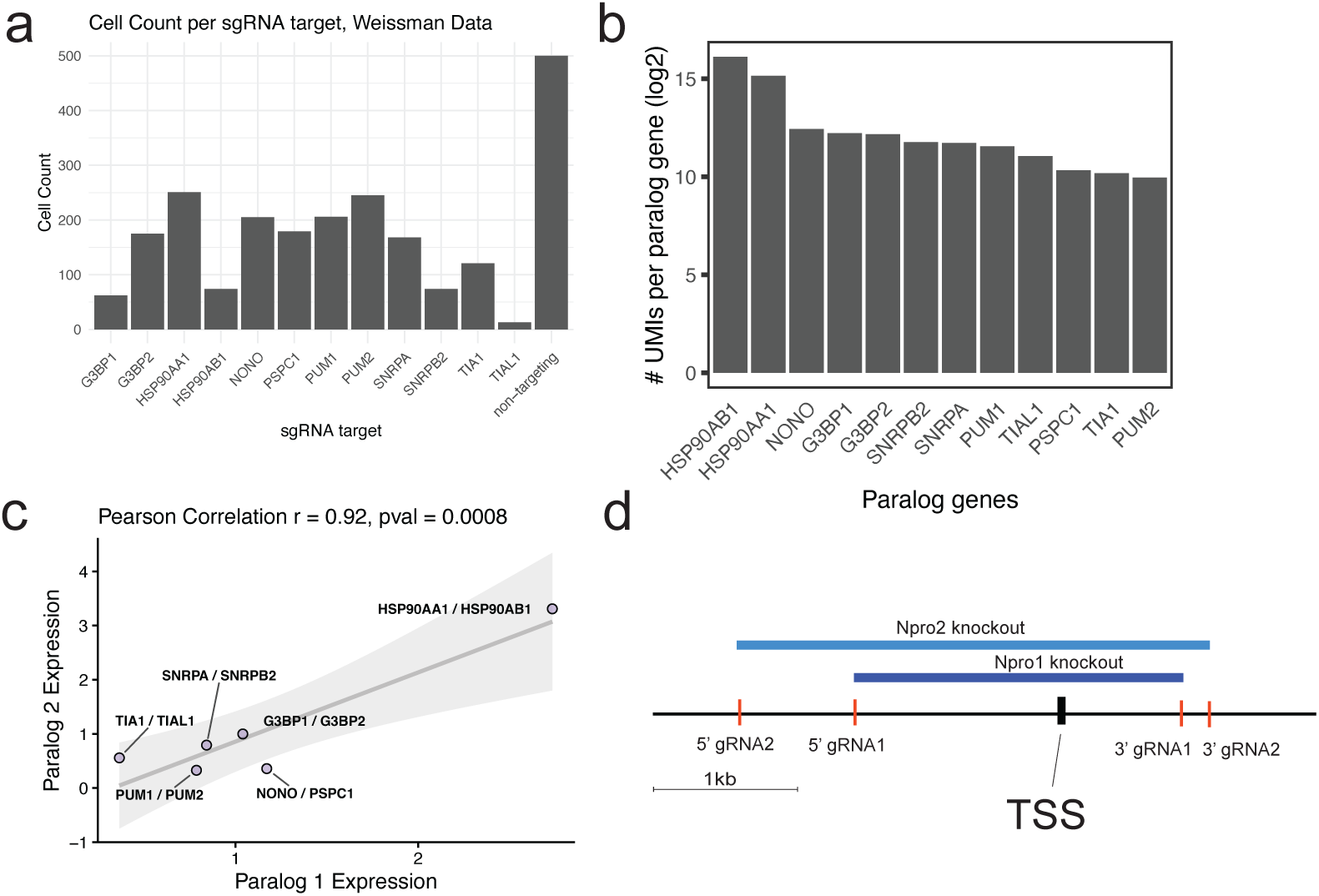
a) Cell counts per sgRNA target in Replogle dataset. b) UMI counts per paralog gene assessed in Replogle *et al* dataset. c) Correlation of paralog pair expression in Replogle *et al* dataset d) Schematic of pgRNAs targeting *NONO* promoter locus.

**Supplementary Figure 4.**
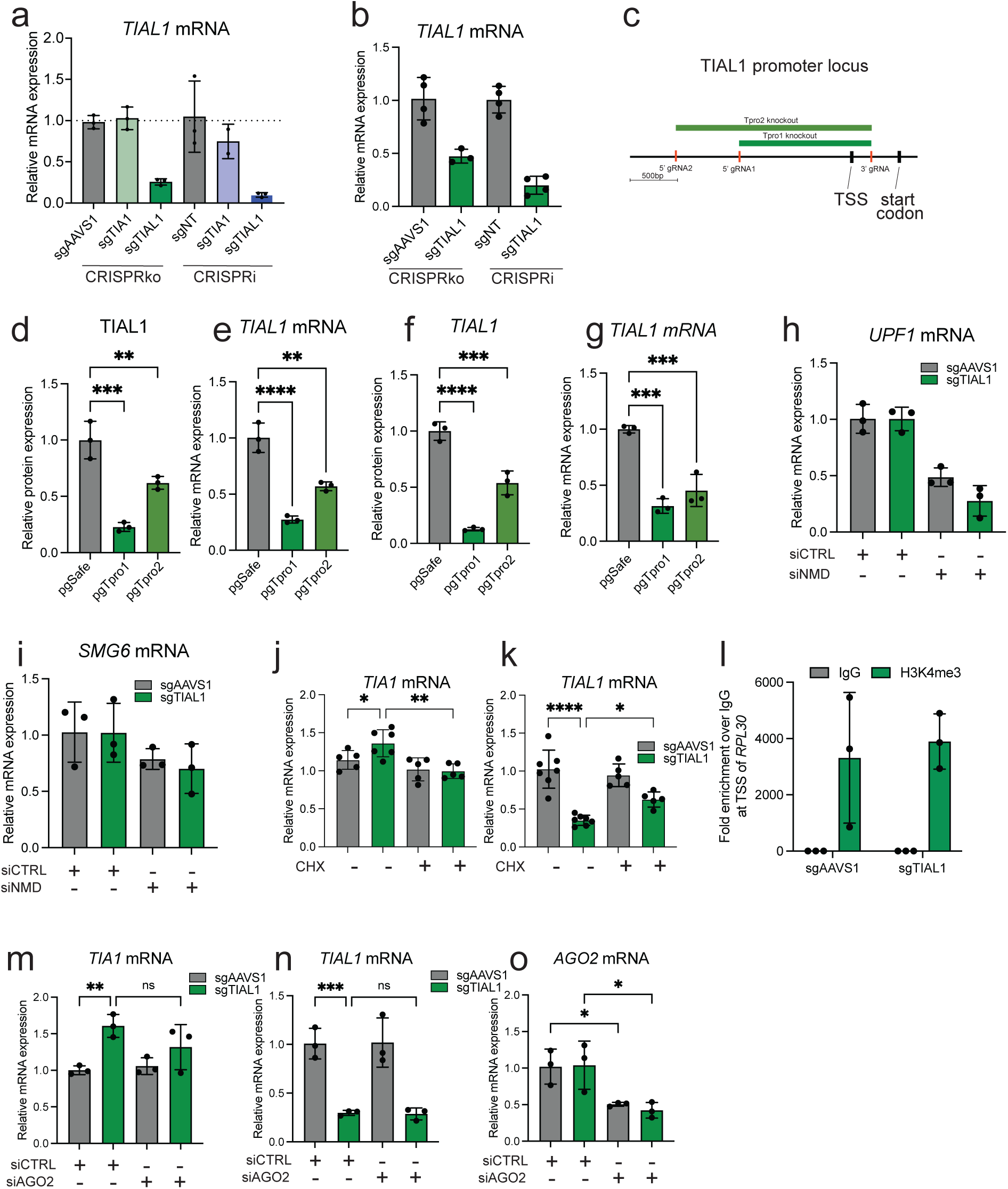
a) *TIAL1* mRNA expression following treatment with indicated gRNAs in PC9 cells. b) *TIAL1* mRNA expression following treatment with indicated gRNAs in Calu6 cells. c) Schematic of pgRNAs targeting *TIAL1* promoter locus. d) TIAL1 protein expression in PC9 cells following treatment with pgRNAs targeting promoter locus, three independent replicates, one-way ANOVA with Sidak’s multiple comparisons test, **p<0.01, ***p<0.001. e) *TIAL1* mRNA expression in PC9 cells following treatment with pgRNAs targeting promoter locus, three independent replicates, one-way ANOVA with Sidak’s multiple comparisons test, **p<0.01, ****p<0.0001. f) TIAL1 protein expression in Calu6 cells following treatment with pgRNAs targeting promoter locus, three independent replicates, one-way ANOVA with Sidak’s multiple comparisons test, **p<0.01, ***p<0.001. g) *TIAL1* mRNA expression in Calu6 cells following treatment with pgRNAs targeting promoter locus, three independent replicates, one-way ANOVA with Sidak’s multiple comparisons test, **p<0.01, ****p<0.0001. h) *TIA1* mRNA expression following treatment of PC9 cells expressing indicated gRNAs with cycloheximide at 100mg/mL for 8hours. Five independent replicates, one-way ANOVA with Sidak’s multiple comparisons test. *p<0.05, **p<0.01. i) *TIAL1* mRNA expression following treatment of PC9 cells expressing indicated gRNAs with cycloheximide at 100mg/mL for 8hours. Five independent replicates, one-way ANOVA with Sidak’s multiple comparisons test. *p<0.05, ****p<0.0001. j) *UPF1* expression in PC9 cells following treatment with indicated siRNAs for 72h, three independent replicates. k) *SMG6* expression in PC9 cells following treatment with indicated siRNAs for 72h, three independent replicates. l) Fold enrichment of activating histone mark H3K4me3 compared to IgG at promoter of control gene RPL30 following treatment with control sgAAVS1 or sgTIAL1 sgRNAs in PC9 cells. Three independent replicates. m) *TIA1* mRNA expression in PC9 cells following treatment with indicated siRNAs for 72h, three independent replicates, one-way ANOVA with Sidak’s multiple comparisons test, **p<0.01 n) *TIAL1* mRNA expression in PC9 cells following treatment with indicated siRNAs for 72h, three independent replicates, one-way ANOVA with Sidak’s multiple comparisons test, ***p<0.001 o) *AGO2* mRNA expression in PC9 cells following treatment with indicated siRNAs for 72h, three independent replicates, one-way ANOVA with Sidak’s multiple comparisons test, *p<0.01

